# Computational Elucidation of Recombinant Fusion Protein Effect on Peptide-Directed Nanoparticles

**DOI:** 10.1101/2021.03.11.432607

**Authors:** Adithya Polasa, Imann Mosleh, James Losey, Alireza Abbaspourrad, Robert Beitle, Mahmoud Moradi

## Abstract

Nanoparticles synthesized using various peptides have optimized properties and functional abilities which can be achieved via peptide flexibility and site specificity. Using peptide Pd4 and other alanine substitution combinations of Pd4 attached to a green fluorescent protein (GFPuv), nanoparticles with well-defined sizes that are soluble in aqueous solutions can be produced. In this study, extensive molecular dynamics simulations explored the structural and functional differences between the free peptides and the peptides bound to the GFPuv used in nanoparticle production. Binding affinities of histidines of Pd4 peptide and its two mutants A6 and A11 to a palladium atom were calculated using the free energy perturbation method. Interestingly, the average particle sizes obtained from transmission electron microscopy (TEM) images correlated with our calculated free energies of different peptide sequences. Remarkably, when the peptide was bound to GFPuv, the free energies of histidine were very similar in the wild-type and other mutated peptides. However, this trend is not observed with free peptide simulations, where binding affinities differ by mutation of histidine residues. This study describes, at a molecular level, the role of amino acid sequence on binding affinity of the peptide to the surface of the palladium particles, and the functional ability of the GFPuv protein controlling these free energies irrespective of peptide sequence. Our study will provide a framework for designing free and protein attached peptides that facilitate peptide-mediated nanoparticle formation with well-regulated properties.

## Introduction

Nanotechnology is a significant, and empowering innovation with applications to many fields of science and technology that benefit from the control of matter at the atomic scale. ^1^ Research on nanoparticles has generated high enthusiasm because of a wide range of potential applications in the biomedical, agricultural, optical, and electronic fields. ^1–4^ Advances in science have developed many synthesis and characterization procedures of nanoparticles in the biomedical areas using both inorganic and organic nanoparticles in targeted drug therapy in mammalians. These therapies have shown potential in numerous applications, including anti-tumor treatments that enhance conveyance of restorative operators into tumors.^5^ Many biological processes developed for interactions between nanoparticles and amino acids^6^ could be upgraded based on the peptide sequences and engagement on the inorganic metal surface.^7^ In the last couple of decades many peptides have been introduced to identify inorganic surfaces, ^8–21^ some of which have been utilized to produce inorganic materials.^20–22^ Production of nanoparticles with different size and shape and aggregation stability using peptide immobilization is useful in sub-areas of biotechnology such as catalysis,^23^ sensors, ^24–26^ and bioanalytical procedures.^27–29^

Changes to the amino acid sequence of a peptide used in the synthesis of a nanoparticle can alter the surface properties of the nanoparticles that are produced. Studies have found evidence that oligopeptides with tryptophan and tyrosine in their sequence are potentially involved in reducing metal ions into their respective metals forming nanoparticles.^30–32^ In contrast, histidine (His) containing oligopeptides, bound to the material surfaces experimentally, have increased the open interaction sites between the solvent and metallic surface. ^33^ Recent computational simulations have demonstrated that aromatic residues of His10 and His12 for Pd2 and His6 and His11 for Pd4 have a lower surface interaction and mobility on the surface of palladium nanoparticles compared to other peptides in the study. ^33^ Substitution of these His residues with an alanine (Ala) has affected the reactivity and nanoparticle fabrication capability of the peptide, resulting in varying turn over frequencies (TOFs) for the Stille coupling reaction. ^33^ Furthermore, a minor mutation in the amino acid sequence, cysteine and alanine modifications, can drastically attenuate the surface structure of nanocatalysts such that absorption energy of peptide to the palladium nanoparticle decreases.^34,35^ The difference between the His6 and His11 in the Pd4 peptide has been the main focus of some studies^7,34^ as mutations of any of these histidines have completely modified the structural and functional abilities of the peptide. A computational study has proposed that His6 of a peptide PD4 has a slightly lower surface interaction energy and mobility of the residue on the surface of the palladium substrate than His11. ^33^ However, His11 has more interaction sites than His6, which explains the contradiction in the free energy profile of these residues in many studies. ^7,33,34^

Control of nanoparticle particle has also been achieved in fusion peptides. Attaching a fluorescent protein into the structure of a peptide provided much more control over the size, fabrication, and delivery of the nanoparticle. ^36^ Additionally, a green fluorescent protein (GFPuv) not only helped in monitoring the synthesis of the nanoparticle but also resulted in synthesizing uniform nanocatalysts in a single-step process. ^37,38^

This study seeks to better understand how the structure and function of the free peptide changes with altered histidines in the amino acid sequence through the use of computer simulations. The subjects of this molecular dynamics (MD) simulation study are the peptide models Pd4 (TSNAV**H**PTLR**H**L) and alanine mutants substitution called A6 (TSNAV**A**PTLR**H**L) and A11 (TSNAV**H**PTLR**A**L) (Fig. 1A-C). Additionally, for each peptide described above, computationally generated GFPuv bound peptide models (Fig. 1D-F) were created. Following structural modeling of the peptides, MD simulations were performed in an aqueous environment, and free energy calculations were done to measure the absolute binding free energy of His and palladium (Pd) ion extensively. Experimental GFPuv mediated palladium nanoparticle synthesis results for particle size and TOF were gathered to validate our results and provide a comparison to existing experimental data for free peptides.

**Fig. 1.**
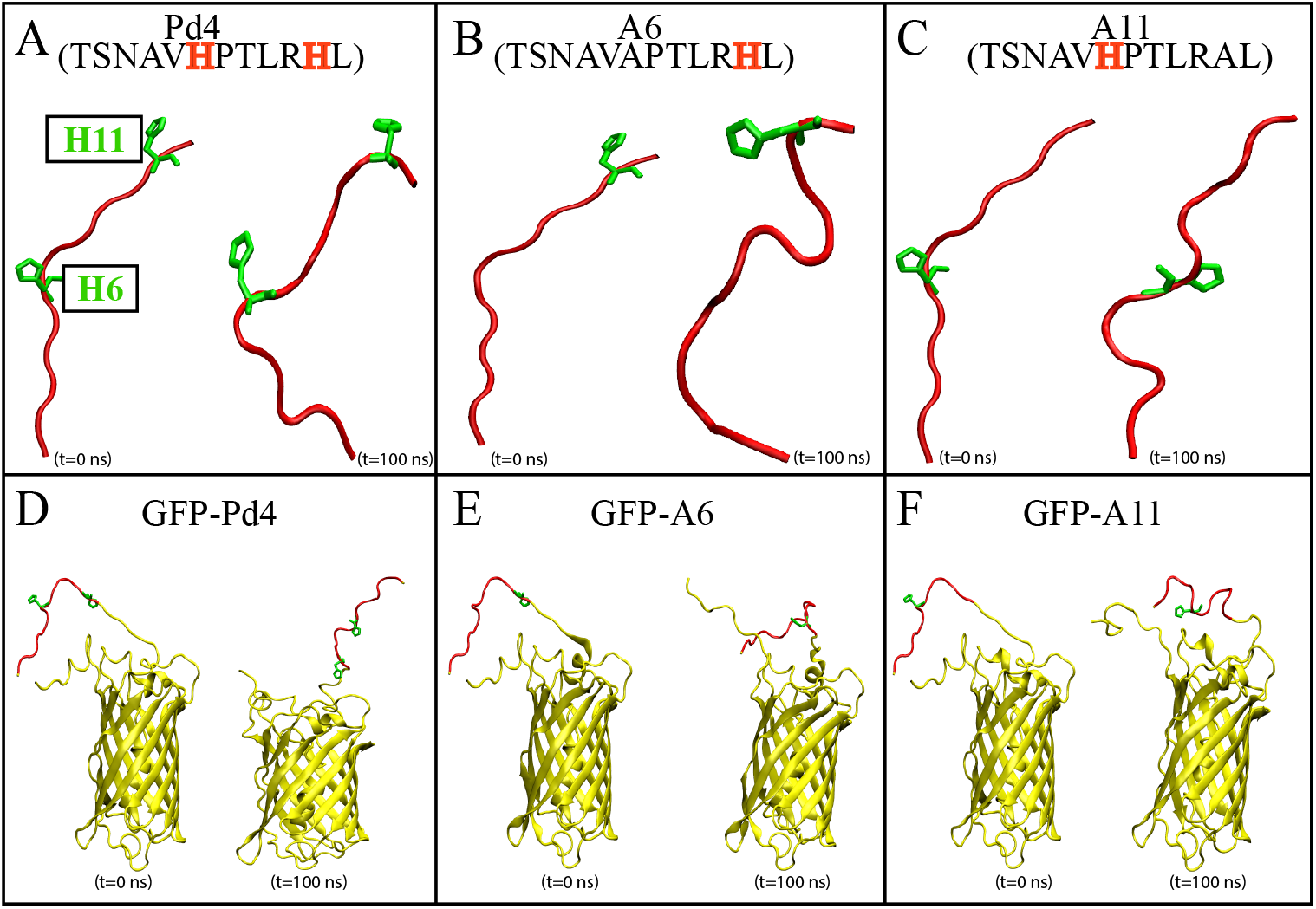
The initial and final MD snapshots of Pd4, A6, A11 (A-C), GFP-Pd4, GFP-A6 and GFP-A11 (D-F). The peptide in each protein is colored red and GFPuv is colored yellow (D-F). Histidines in the peptide sequence are colored green.

These simulations added detail to the the sequence-dependent structural and functional dynamics of the peptides at the atomic level. The results of this study provide insight for designing new oligopeptides and for understanding the production of biologically applicable nanoparticles.

## Methods

All-atom MD simulations were performed to obtain a relaxed structure of peptide models along with conformational dynamics and binding energies of His residues on the surface of Pd atoms. For the fusion peptides, the crystal structure of GFPuv (PDB: 1W7S) ^39^ was obtained from RCSB.org with a resolution of 1.85 Å. Initially, Modeller^40^ homology modelling software was used for the construction of all peptide and GFPuv fusions (Fig. 1). Next, CHARMM-GUI ^41,42^ web-server was used to build the MD simulation models of peptides and GFPuv fusions in aqueous solution of TIP3P^43^ water. The addition of 0.15mM NaCl to every system neutralized the electrostatic charge. The total number of atoms for peptide and GFPuv fusion peptide systems was ≈22300 and ≈68000, respectively. NAMD 2.13^44^ was utilized to run the MD simulations with periodic boundary conditions (PBC) at 310K in the NPT ensemble, and 1 atm pressure was maintained using the Nosé-Hoover Langevin piston method.^45,46^ Every system was equilibrated for 100 nanoseconds (ns) using CHARMM36 all-atom force field parameters. The binding free energy for each peptide was then calculated using the alchemical free-energy perturbation (FEP) procedure. This procedure was repeated on 20 different conformations of each peptide resulted from equilibrium simulations.

### Fusion Protein Preparation

According to the codon preference of *E.coli*, plasmids encoding Pd4, A6, and A11 peptides fused to GFPuv were constructed. The fragment containing codons of peptides were introduced to 5 end of the GFPuv gene using primer forward primers through polymerase chain reaction (PCR). Synthetic genes containing desired peptides and addgene-plasmid-51559 were double digested with EcoRI and XbaI restriction enzymes before the ligation. Finally, the designed DNA containing desired peptides were ligated to the DNA to construct plasmids containing GFPuv fusion proteins. Bacterial lysates from arabinose induced cells were obtained, followed a method adapted from our previous research.^38^ Protein concentrations were determined using the DC Protein Assay (Bio-Rad, Hercules, CA).

### Nanoparticle synthesis

To synthesize palladium nanoparticles (Pd NPs) at room temperature, 0.16 mg K_2_PdCl_4_ was added to synthesis mixtures (one milliliter total volume) containing 0.23 mg fusion protein. These amounts result in a 2:1 ratio of Pd^+2^ to Pd4. After mixing for 0.5 hour, 1.5 mg NaBH_4_ was added to the mixture to reduce Pd^+2^ ions to metalic Pd and NPs were formed rapidly after reduction indicated by a color change (yellow to light brown). The NP shape and size distribution were analyzed using TEM. A droplet containing ten microliter samples of the reaction mixture were placed on a 300 mesh standard lacey carbon grid. Measurements with an FEI Titan 80–300 determined NP size distribution and morphology. Finally, TEM images were analyzed using ImageJ software.^38^

### Screening of reaction parameters for Suzuki-Miyaura coupling reaction

For optimizing the reaction condition, a method developed by Mosleh *et al.,* was adopted.^38^ The different reaction conditions including base, temperature, and solvent were evaluated using the model coupling reaction (Table S1). The reaction did not result in high yield when KOtBu, K_2_HPO_4_, and KH_2_PO_4_ (entries 1-3) were used as base while the reaction proceeded with excellent catalytic activity in the presence of K_2_CO_3_ (entry 4). The catalytic performance of reaction under different temperatures indicated that by increasing the temperature, higher yields could be obtained. Indeed, the presence of EtOH in water-contained solvents as a green solvent with the ratio of 1:1 was found to be the best solvent for Suzuki-Miyaura coupling reaction (entries 6-8).

### Screening of reaction parameters for Stille coupling reaction

To obtain the optimized condition for Stille coupling reaction, the reaction of iodobenzene and phenyltin trichloride was used as the model reaction and other reaction conditions including base, temperature, and catalyst loading were evaluated (Table S2). For the base study, a series of bases were explored and K_3_PO_4_ was found to be the best base as the yield of biphenyl production was 97% (entries 1-4). Higher yields could be obtained when higher temperatures were used. Although 70% yield of biphenyl was proceeded at 60 °C, the reaction was performed for 20 hrs. Employing 80 °C resulted in biphenyl production with 96% yield after 6 hrs. Furthermore, increasing the amount of catalyst did not alter the yield of biphenyl while lower catalytic activities were observed when 2 mmol% and 1 mmol% of Pd was present in the reaction (entries 7-9).

### Free energy calculations

The binding free energy of His in peptide region and palladium atom for all systems mentioned above were calculated (Fig. 1). All simulations were carried out using NAMD 2.13^44^ and TIP3P^43^ water. For FEP calculations, 20 snapshots for 100 ns equilibrated system were taken and simulations were performed changing *λ* (through eventually decoupling outgoing atom interactions and coupling incoming atom interactions) from zero to one or from one to zero, respectively. Hydration free energy of palladium ion was carried out under PBC conditions at constant pressure. The systems were simulated with a 2 fs time step using Langevin dynamics at a temperature of 310K.

In all calculations, parsefep plugin^47^ calculated the free energies for the equilibrated simulations. 20 independent FEP calculations were performed consecutively for each system for binding calculation. To examine the binding free energy quantitatively, the thermodynamic cycle illustrated in the Fig. S5 was constructed. This cycle connects the binding of palladium atom with the histidine in peptide individually for each system from unbound state to bound state, i.e., Δ*G*_*unbound.→bound*_= Δ*G*_2_ with solvation of palladium atom in aqueous solution Δ*G*_*vacuum.→aqueous*_= Δ*G*_1_. Overall, the binding free energy of palladium was calculated based on the equation Δ*G*_*binding*_ = Δ*G*_1_ - Δ*G*_2_ derived from the thermodynamic cycle proposed (Fig. S5).

## Results and Discussion

### Histidine-palladium binding free energies for free peptides

The binding free energies of the histidines with the palladium atom are reported in Table 1 along with average particle size and TOF from Ref. 7 and Ref. 34. The difference between the binding free energies of histidine to the palladium atom shows the importance of the histidine in the sequence of the free peptide. For systems with His11 present, the free energy of binding with His11 was −156.86±33.97(kcal/mol) and −145.29±31.08(kcal/mol) for Pd4 and A6, respectively. On the other hand, the free energy for binding with His6, in Pd4 and A11, were −89.00±24.74(kcal/mol) and −118.33±26.46(kcal/mol), greater than His11 binding free energies. The lower free energy for binding with His11 made it the preferred palladium binding site when it was available, which directly relates to its high affinity for binding with the palladium nanoparticle.

**Table 1:** Binding free energy of Pd nanoparticles with free peptide.

| Peptide | Binding Free Energy<br>(kcal/mol) | | | TOF<br>($h^{-1}$) | | Average Particle<br>Size (nm) | |
| --- | --- | --- | --- | --- | --- | --- | --- |
|  | His6 | His11 | Effective | Ref. 7 | Ref. 34 | Ref. 7 | Ref. 34 |
| **Pd4** | -89.00±24.74 | -156.86±33.97 | -156.86 | 2234±99 | 2200±100 | 2.1±0.4 | 1.9±0.3 |
| **A6** | | -145.29±31.08 | -145.29 | 5224±381 | 5200±400 | 2.2±0.7 | 2.2±0.4 |
| **A11** | -118.33±26.46 | | -118.33 | 1298±107 | 1300±10 | 2.6±0.4 | 2.4±0.5 |

The binding free energies from Table 1 provided insight into existing catalytic rate data for TOF from Ref. 7 and Ref. 34. When only the His11 was present, as in A6, the binding free energy increased a small amount compared with Pd4, with overlapping error bars. The TOF for A6 increased more than double, from 2200±100 h^−1^ to 5200±400 h^−1^. Conversely, when only the His6 is present, as in A11, the TOF decreased to 1298±107 *h*^−1^. In Pd4, both histidine residues were involved in the nanoparticle synthesis reaction. Having all reactivity focused on the His11 resulted in the higher TOF of A6 peptide. With His6 and His11 present in Pd4, the catalytic activity was slowed down because some binding occurred with less reactive His11, despite Pd4 the having lowest binding free energy of the three systems tested.

Nanoparticle size data from TEM measurements displayed a linear relationship with the minimum binding free energy of the histidines in sequence (Fig. 3A). Higher binding free energy of the palladium with the peptide during synthesis correlated with larger particles. The free energy calculations help in understanding the competitive relationships between histidine binding sites that determine catalytic activity of the peptide and the size of the nanoparticle produced. Extensive analysis was conducted on the trajectories of peptide simulation to further explore these differences.

### Secondary structure propensity of residues 6 and 11 of Pd4, A6, and A11

To examine the impact of the histidine mutations on secondary structure in the simulations, the secondary structure of residue 6 and 11 of peptides were analyzed based on *ϕ − ψ* angles plotted on the Ramachandran plot (Fig. 2). This analysis revealed that the *α*_*L*_ propensity of residues was significantly affected by the mutation of His6 and His11. The secondary structure and orientation of the peptide could be a factor for the different catalytic activity.

In Figure 2, the majority of residues were located in the F region of the Ramachandran plot. The mutation of the His6 to an alanine in A6 changed the structure of His11, which features a significantly higher *α*_*L*_ propensity value giving it more *α* helical structure compared to peptides Pd4 and A11. For A11, the mutation of the His11 to an alanine increased the propensity of the *α*_*L*_ secondary structure of residue 11 but not as prominently as in the A6 peptide sequence. This could explain the high reactivity of A11. On the other hand, in the Pd4 peptide *α*_*L*_ propensity of both 6^*th*^ and 11^*th*^ histidines residue have a very close distribution, this trend was not observed in the mutated peptide residues. This secondary structure difference could be a reason for the difference in the TOF and free energy results, since the *α* structure of the amino acid is directly proportional to the reactivity because of the rigid structure of the residue.

**Fig. 2.**
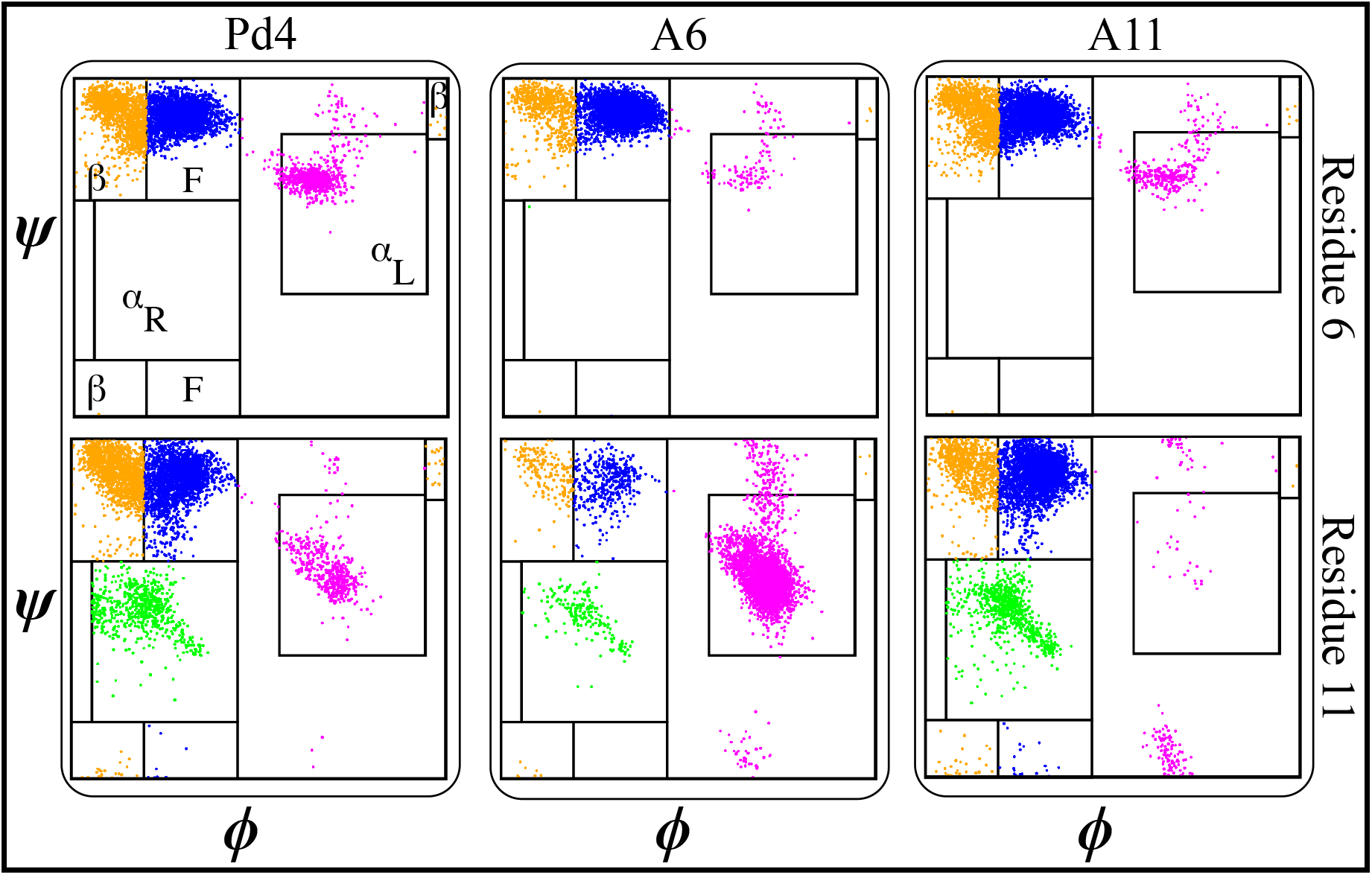
The structural propensity of peptide Pd4, A6, and A11 shown as a Ramachandran plot (x-axis *ϕ* and y-axis *ψ* angles) of residue 6 (top) and residue 11 (bottom). Secondary structures are colored with *α*_*R*_ as green, *α*_*L*_ as magenta, *β* as orange, and F as blue.

### GFPuv fusion peptides simulation and experiment

To study the ability of the GFPuv fusion peptide framework for the production of Pd nanoparticle, equilibrium simulations of peptide bound to the GFPuv protein were conducted to study the structural and functional properties of the peptide region in GFPuv-peptide framework. Palladium nanoparticles were generated *in vitro* using the Stille and Suzuki-Miyaura coupling reactions. A schematic of the different reactions and a plot of the catalytic data is presented in Fig. S1.

The nanoparticles obtained from the three GFPuv fusion constructs were analyzed by TEM to measure particle size (Fig. S6). The binding free energy from simulation is shown with the catalytic results and the TEM particle sizes from the experiment in Table. 2. An average particle sizes of 2.6±0.5(nm), 2.7 ± 0.7(nm), and 2.6±0.4(nm) where observed from nanoparticles prepared from GFPuv fusion Pd4, A6 and A11, respectively. These particle sizes are reasonably close to the linear relationship of observed between binding free energy and particle size. Uniform TOF results were observed for all fusion peptides in Still coupling reactions. Comparing this with Table 1, the TOF of the fusion peptides are higher than Pd4, but lower than the A6 mutant. Results for the Suzuki-Miyaura reactions provided a different TOF value, but the uniform catalytic behavior of the fusion peptides persisted across different reactions.

Kernel density estimation using 100 bootstrap resampling on the TEM particle size results for all GFPuv fusion peptide-mediated nanoparticle synthesis continued the consistent behavior, with a similar density distribution on the kernel map for all peptides (Fig. 3A). The binding free energy, TOF, and average nanoparticle size were very similar for all GFPuv fusion peptides, showing agreement with

**Table 2:** Binding free energy of Pd nanoparticles with GFP fused peptides.

| Peptide | Binding Free Energy (kcal/mol) | | | TOF ($h^{-1}$) | | Average Particle Size (nm) |
| --- | --- | --- | --- | --- | --- | --- |
|  | His6 | His11 | Effective | Stille | Suzuki-Miyaura |  |
| **GFP-Pd4** | -118.04±26.71 | -122.28±27.24 | -122.28 | 2945.33±177.58 | 11731±1452.58 | 2.6±0.5 |
| **GFP-A6** | | -121.62±26.99 | -121.6 | 2912±173.6 | 11093±8333.06 | 2.7±0.7 |
| **GFP-A11** | -118.17±27.06 | | -118.17 | 2942±113.01 | 10867.33±767.6 | 2.6±0.4 |

**Fig. 3.**
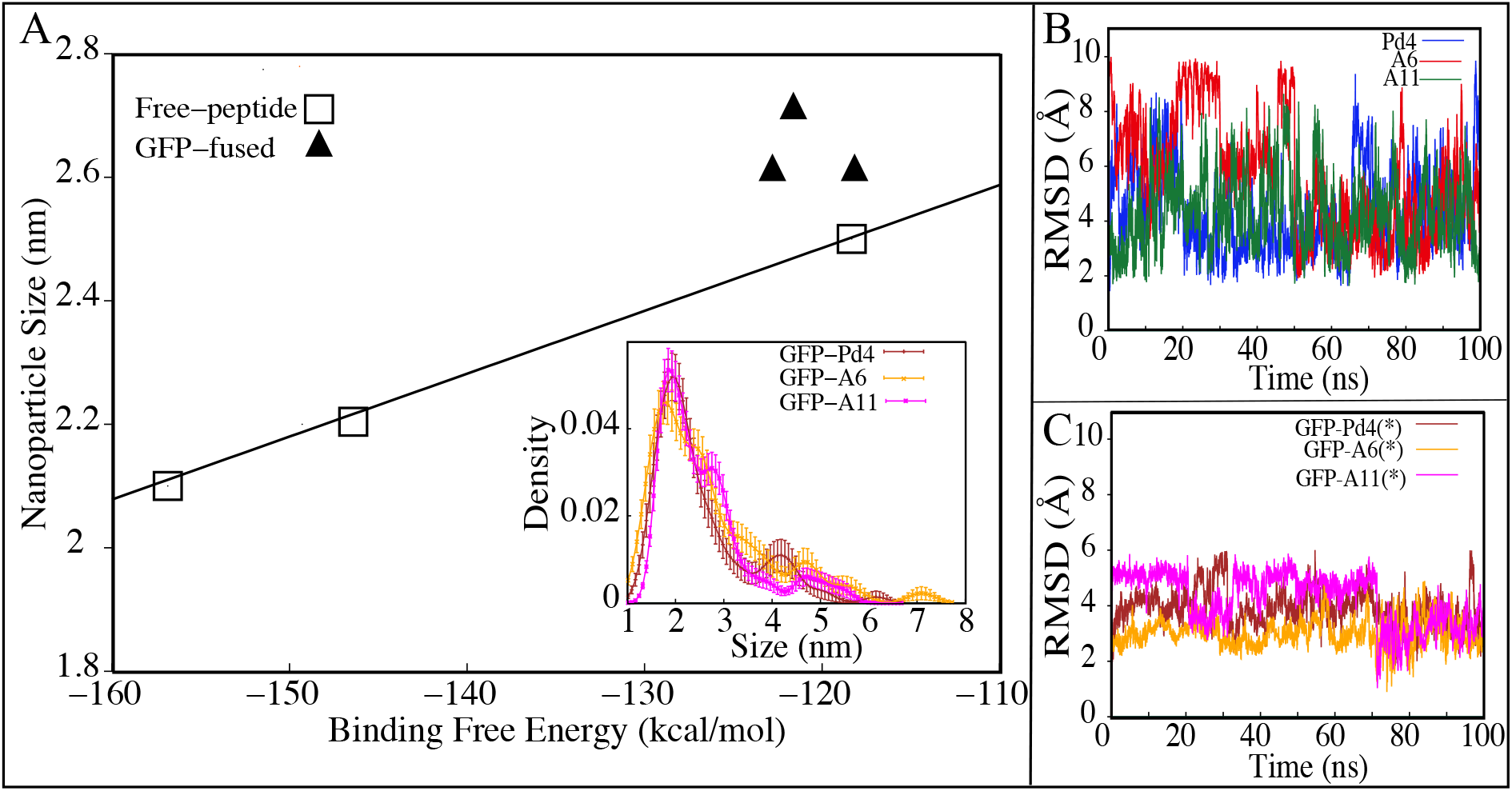
(A) Nanoparticle size as a function of binding free energies (line represents linear regression of free peptide results). The insert shows the kernel density estimation of 100 bootstrap resamples of experimentally measured nanoparticle sizes. Nanoparticle size distribution synthesised using GFPuv fusion peptides. (B-C) RMSD profile of free peptides and GFPuv fused peptides, respectively. (*) Only the peptide region was used for RMSD calculation in (C).

The root mean square deviation (RMSD) of the peptide portion in the GFP bound simulations were stable in all cases in our simulations (Fig. 3C), which suggests that GFPuv stabilizes the entire peptide conformation in nanoparticle preparation. The RMSD of free peptide backbones fluctuated higher than the RMSD of the bound state (Fig. 3B). From the RMSD results one might postulate that conformational stability of protein/peptide was important in synthesizing uniform nanocatalysts.

Previous studies ^6,33^ of nanoparticle synthesis using free peptides produced nanoparticles with a slightly varying particle size/width distribution indicating that the effective binding free energy between the histidine and Pd impacted the nanoparticle growth distribution in the free peptide method. In free peptide nanomaterial synthesis, a slight distribution error was observed in the nanomaterial size with replacement of histidine in Pd4. On the other hand, mutation of the histidine has not affected any of our GFPuv fusion peptide experiments. The bound GFPuv controlled the structural fluctuations of all the peptides tested during nanoparticle production.

### Implications of GFPuv on structural propensity of peptide and TOF

A secondary structure analysis of the GFPuv fusion peptides simulations found further differences with the free peptide simulations. The Ramachandran plots of peptides Pd4, A6, and A11 bound to GFPuv are shown in Figure 4. The coloring cluster of the plots are consistent with Figure 2. The propensity for *α*_*L*_ states was lower in the all the bound peptide simulations. Also, the relationship between histidine and *α*_*L*_ propensity observed in the free peptide simulations did not exist for the GFPuv fusion bound peptides. The presence of histidine at position 6 and 11 had either no correlation or inverse correlation to the *α*_*L*_ propensity. For GFP-Pd4, the *α*_*L*_ propensity was similar for residue 6 and 11,but in GFP-A6 simulations, the histidine at residue 6 showed the increase in *α*_*L*_ propensity, while residue 11 had a lower *α*_*L*_ propensity. Finally, GFP-A11 showed near zero *α*_*L*_ states on either residue 6 or 11, regardless of the presence of a histidine.

**Fig. 4.**
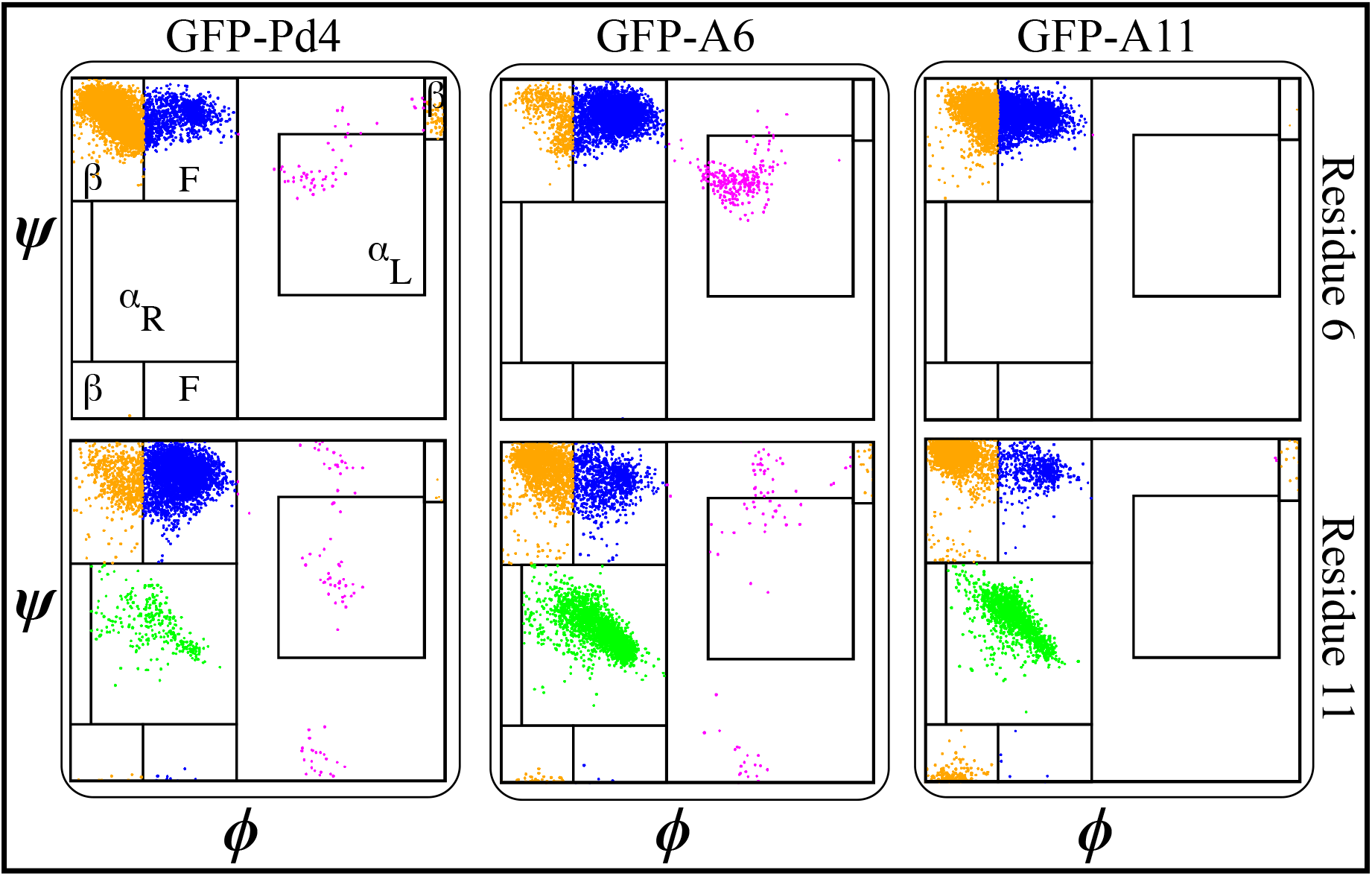
Structural propensity of the GFPuv fused peptide Pd4, A6, and A11 shown as Ramachandran plots (x-axis *ϕ* and y-axis *ψ* angles) of residue 6 (top row) and residue 11 (bottom row). Secondary structures are colored with *α*_*R*_ as green, *α*_*L*_ as magenta, *β* as orange, and F as blue.

### Hydrogen bond analysis

Hydrogen bonds are significant interactions in proteins and peptides, contributing to backbone conformational stability differences. To quantify hydrogen bonds in the peptide backbones, bond length and angle cutoffs of 3.0Å and 30°, respectively, were applied to the simulated trajectories. In the hydrogen bond analysis in Figure 5A, the A6 peptide formed more stabilized backbone hydrogen bond interactions over the length of the simulation compared with Pd4 and A11, which have mostly resulted in zero hydrogen bonds in occupancy percentage.

**Fig. 5.**
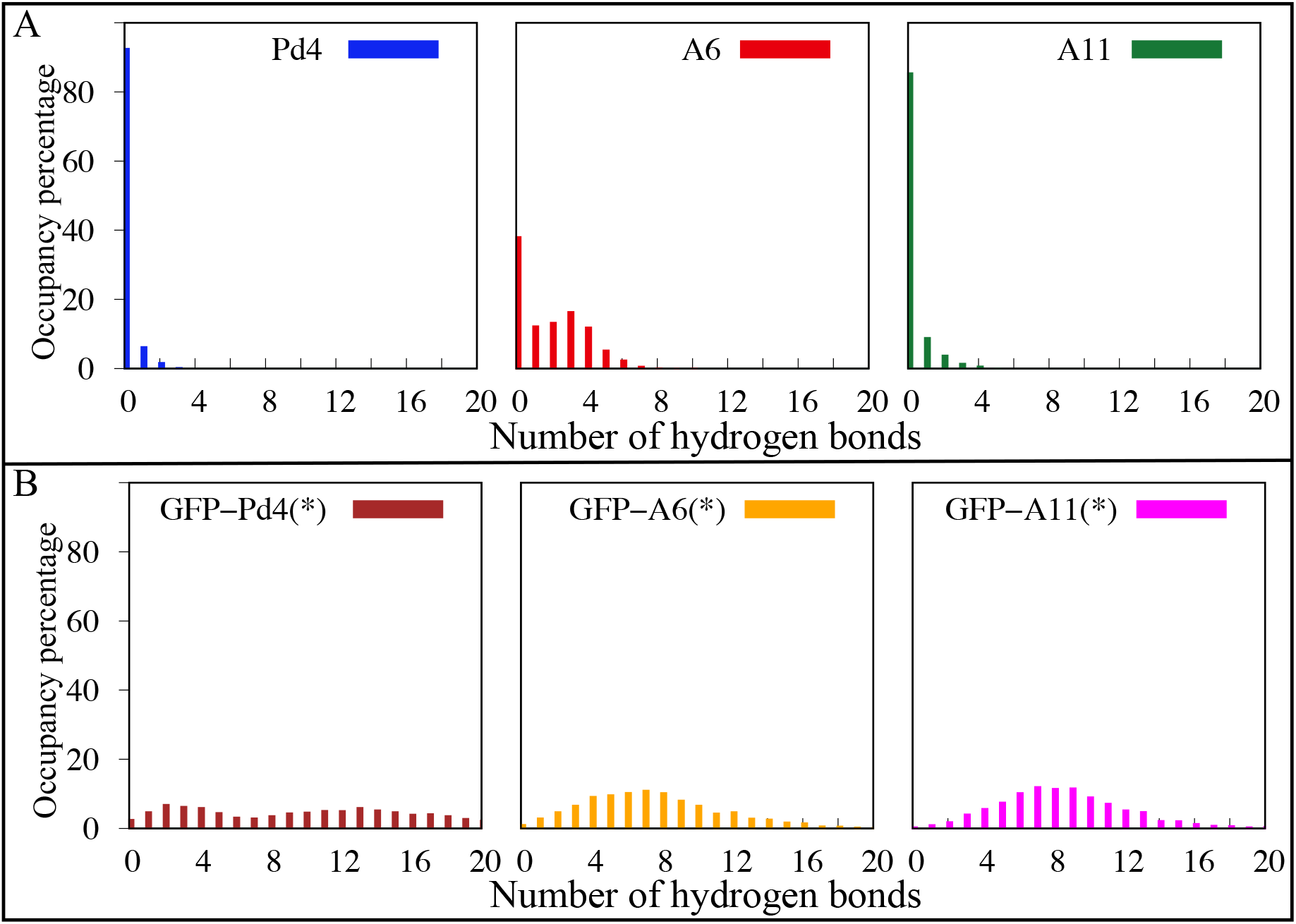
The occupancy percentage of hydrogen bonds calculated from the MD simulations for the (A) free peptides and (B) GFPuv fused peptide region. (*) Only the peptide region in the GFPuv fused peptide simulations was used for hydrogen bond analysis.

In Fig. 5B, hydrogen bond interactions of just the peptide region for GFPuv fusion peptides are reported over 100 ns of equilibrium simulation time. The distribution of hydrogen bonds was qualitatively similar for all the GFPuv fusion peptides, with a broad distribution of number of hydrogen bonds. In the free peptide simulations, the A6 mutation contained the greatest *α*_*L*_ propensity, and also the most persistent hydrogen bonding. The presence of GFPuv stabilized the RMSD of the peptides without increasing the *α*_*L*_ propensity, seen in Fig. 3C.

Specific hydrogen bonds that occurred in the free peptide simulations were identified. Two interactions involving mutation (R10-P7 and A6-L9) have very high occupancy only in A6 mutant peptide. A6-L9 interaction is very strong with an occupancy of 38%, indicating that H6A (A6) might play a key role in conferring protein stability and decreasing binding free energy. Another hydrogen bond R10-P7 (occupancy 46%) was also identified. Both of these hydrogen bonds occurred only in mutant peptide A6.

Principal component analysis (PCA) in Fig. S4 also supports distinct differences in conformations for A6 compared with the qualitatively similar results for Pd4 and A11. Pd4 and A11 had fairly dispersed distributions along PC1 and PC2, but A6 was slightly more compact as stability was increased. From the PCA of the fusion peptides, even more stability was observed, with denser clouds indicating variance was low through the simulations when bound to GFPuv.

## Conclusions

Overall, our computational and experimental results have added explanatory detail to the existing sequence dependent catalytic knowledge of the free Pd4 peptide and Pd nanoparticle. This computational study shows how a single amino acid change in the free peptide sequence can impact the energetic, structural, and catalytic characteristics. Conversely, the GFPuv fusion peptides showed little response to sequence changes in the peptide for free energies and nanoparticle production measures, with the GFPuv influences dominating the sequence mutations.

In the free peptide, the mutation of His6 to alanine in A6 concentrated palladium reactions on residue 11, where binding free energy was most favorable. Also, the His6 mutation led to the increased *α*_*L*_ propensity of residue 11 and the hydrogen bond occupation in the peptide backbone, stabilizing the peptide.

GFPuv acted as a stabilizer when bound to the peptides, with similar effective free energy minima, for wild-type and mutant peptides. Palladium binding free energy at the histidines was moderated, but the entire peptide was stabilized when bound, exhibiting similar behavior for peptide RMSD, secondary structure, and hydrogen bond occupation. The uniformity in experimental TOF and particle size results was consistent with the stability observations from simulation, showing consistent behavior from all peptides. The lower binding free energy led to lower Stille coupling TOF when compared to the free A6 peptide, though TOF was greater than free peptides with lower effective binding free energy. This indicates that binding free energy accounted for only part of the catalytic behavior in this system. With the lower binding free energy, the average size of the nanoparticles increased following the approximately linear behavior shown in Figure 3A.

This research demonstrates a new study for designing multi-functional peptides with various amino acid domains for the production of nanoparticles from inorganic materials. The approach of this study can be efficiently used in other nanocatalyst production, to explain and potentially identify peptide regions important to nanoparticle production through FEP/MD simulations. With this information, optimization of nanoparticle production reactions would be possible from first principles.

## Supporting information

Supporting Informations

## Acknowledgement

This research is supported by the National Science Foundation under Award CHE-1945465. This research is part of the Blue Waters sustained-petascale computing project, which is supported by the National Science Foundation (awards OCI-0725070 and ACI-1238993) and the state of Illinois. This work also used the Extreme Science and Engineering Discovery Environment (allocation MCB200215), which is supported by National Science Foundation grant number ACI-1548562. This research is also supported by the Arkansas High Performance Computing Center, which is funded through multiple National Science Foundation grants and the Arkansas Economic Development Commission.

## Supporting Information Available

Figures S1–S6 and Tables S1–S2 in Supporting Information provide additional data as discussed in the manuscript.

