## Supporting Informations for "Computational Elucidation of Recombinant Fusion Protein Effect on Peptide-Directed Nanoparticles"

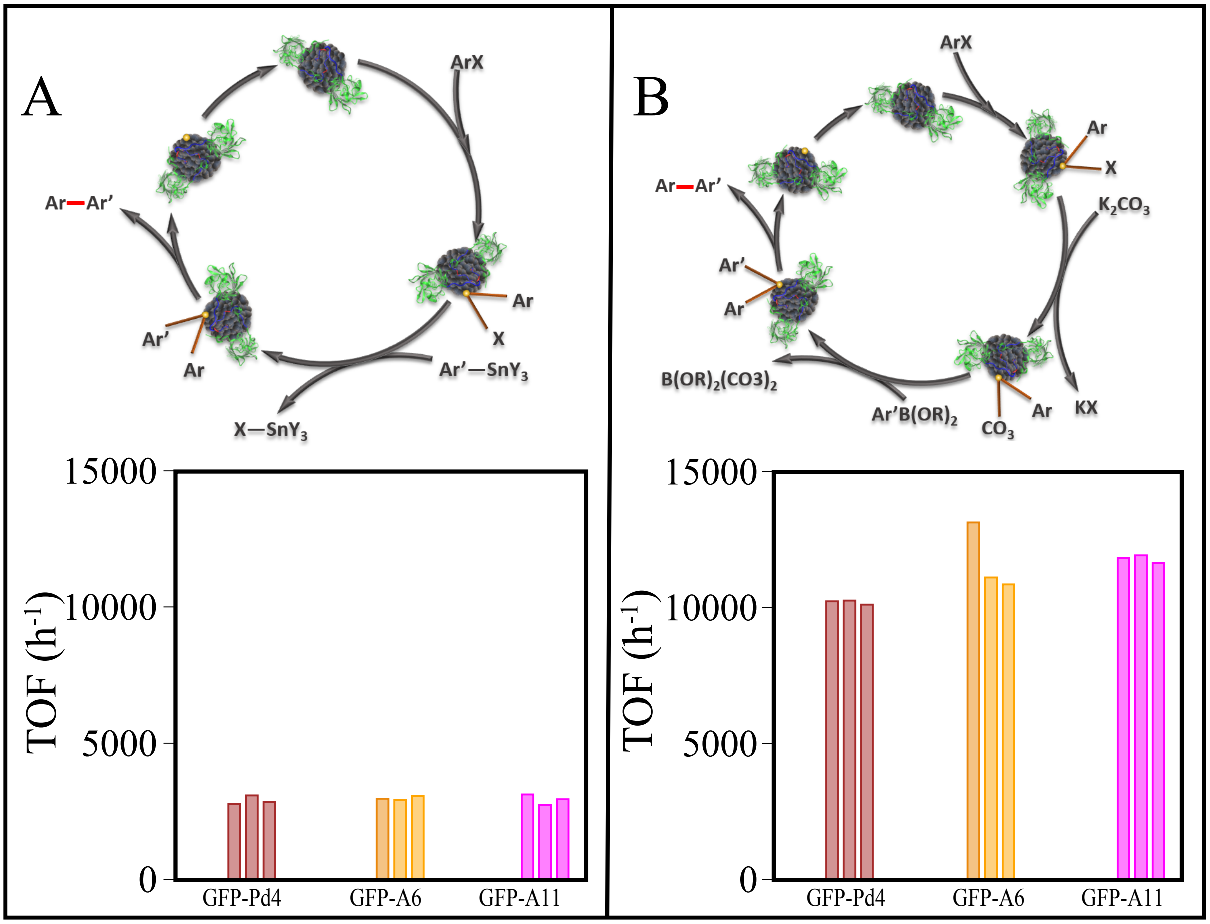


**Fig. S1.** Schematic diagram and TOF of (A) Stille coupling and (B) Suzuki-Miyaura coupling reaction for GFP fused peptides Pd4, A6 and A11 respectively.

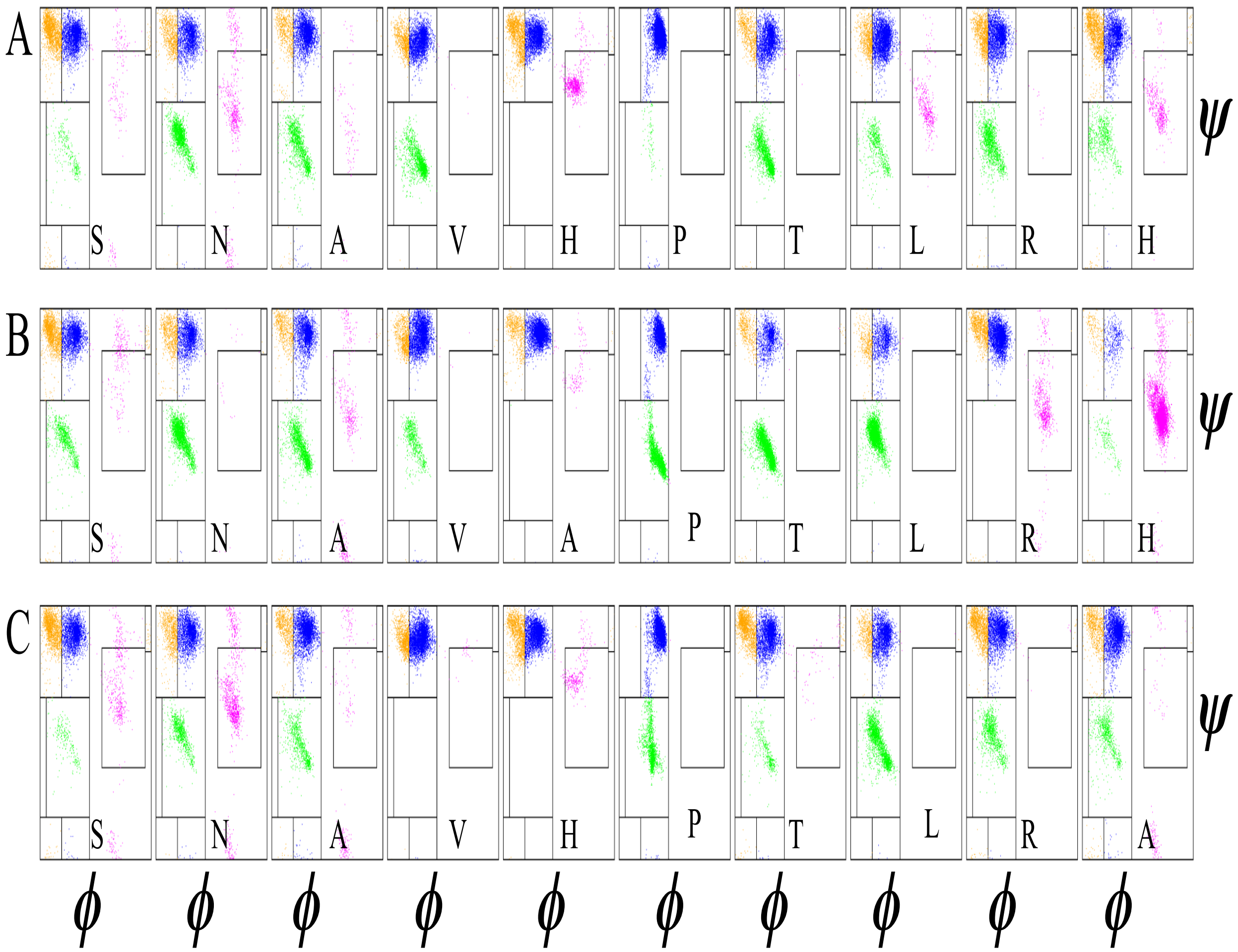


**Fig. S2.** Ramachandran plots of residues in free peptides: (A-C) The backbone structure of the residue in peptides Pd4, A6, and A11 respectively. The region definitions are the same as Figure 1A. Orange, blue, pink, green, and gray clusters identify the $\beta$, F, $\alpha_L$, $\alpha_R$, and N regions, of Ramachandran plot respectively.

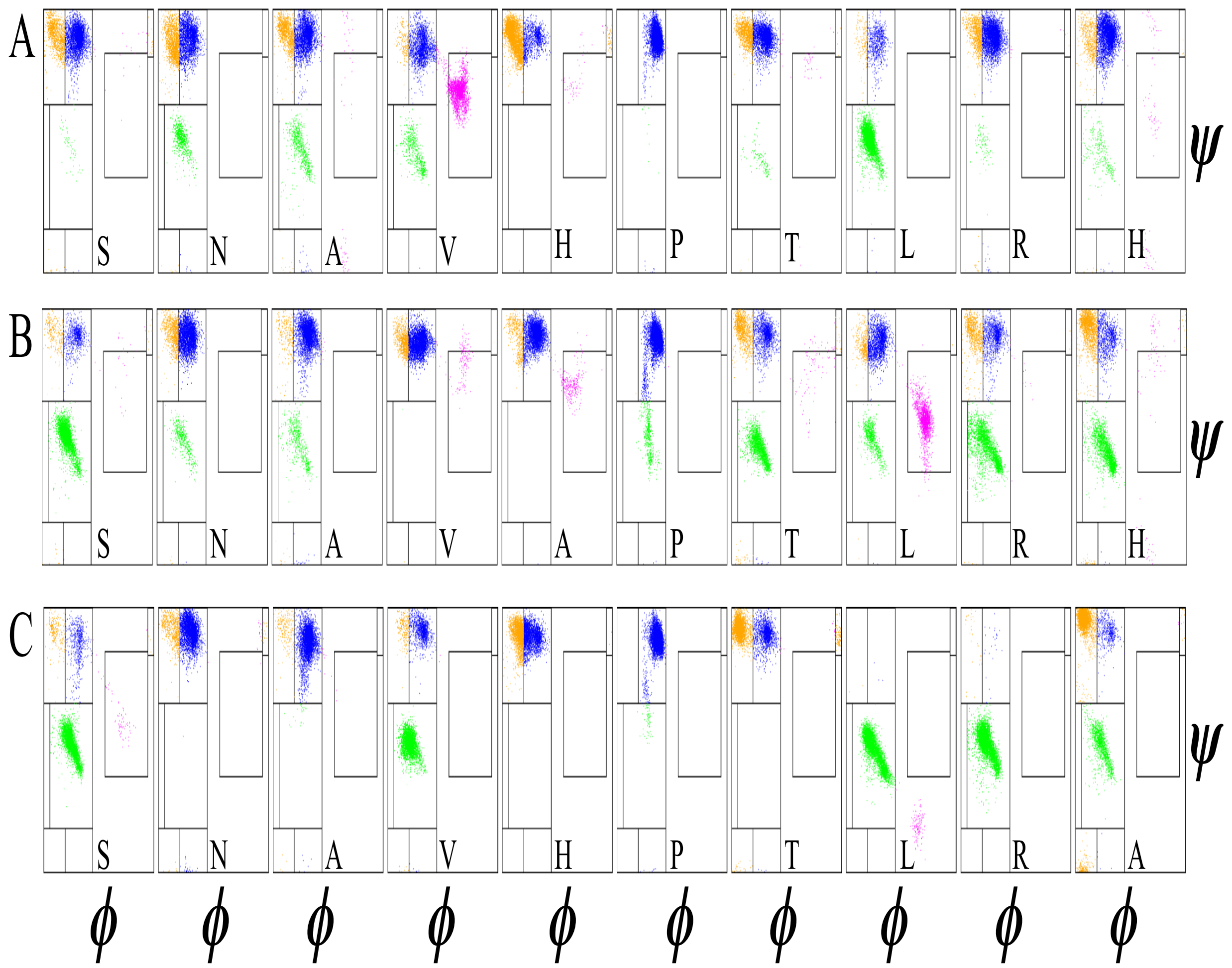


**Fig. S3.** Ramachandran plots of residues in GFPuv fusion peptides: (A-C) The backbone structure of the residue in GFP bound peptides Pd4, A6, and A11 respectively. The region definitions are the same as Figure 1A. Orange, blue, pink, green, and gray clusters identify the $\beta$, F, $\alpha_L$, $\alpha_R$, and N regions, of Ramachandran plot respectively

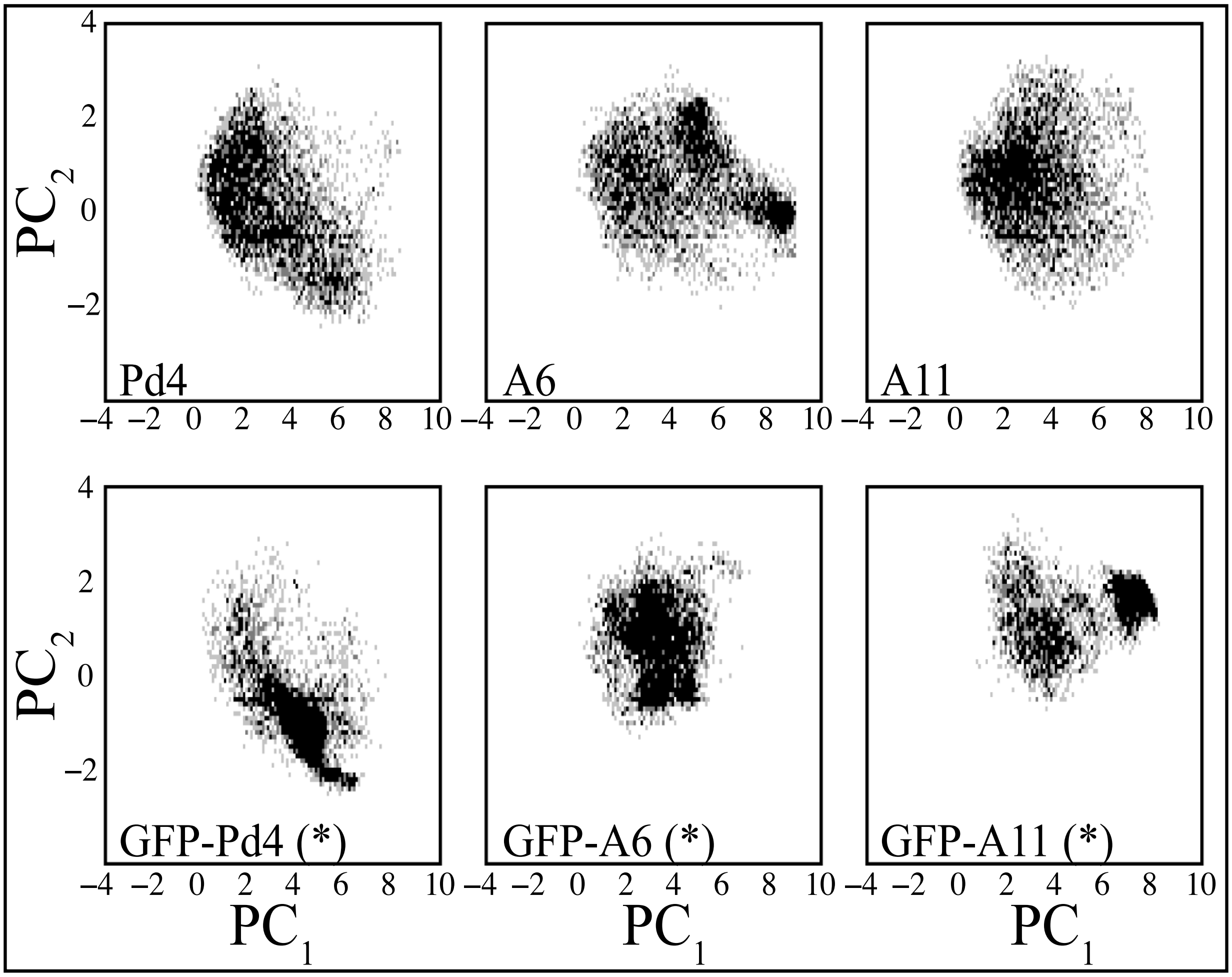


**Fig. S4.** PCA of C-$\alpha$ of peptides in free state and GFP fused simulations. The intensity of the color in the plot represents the relative population of peptide PCA analysis.

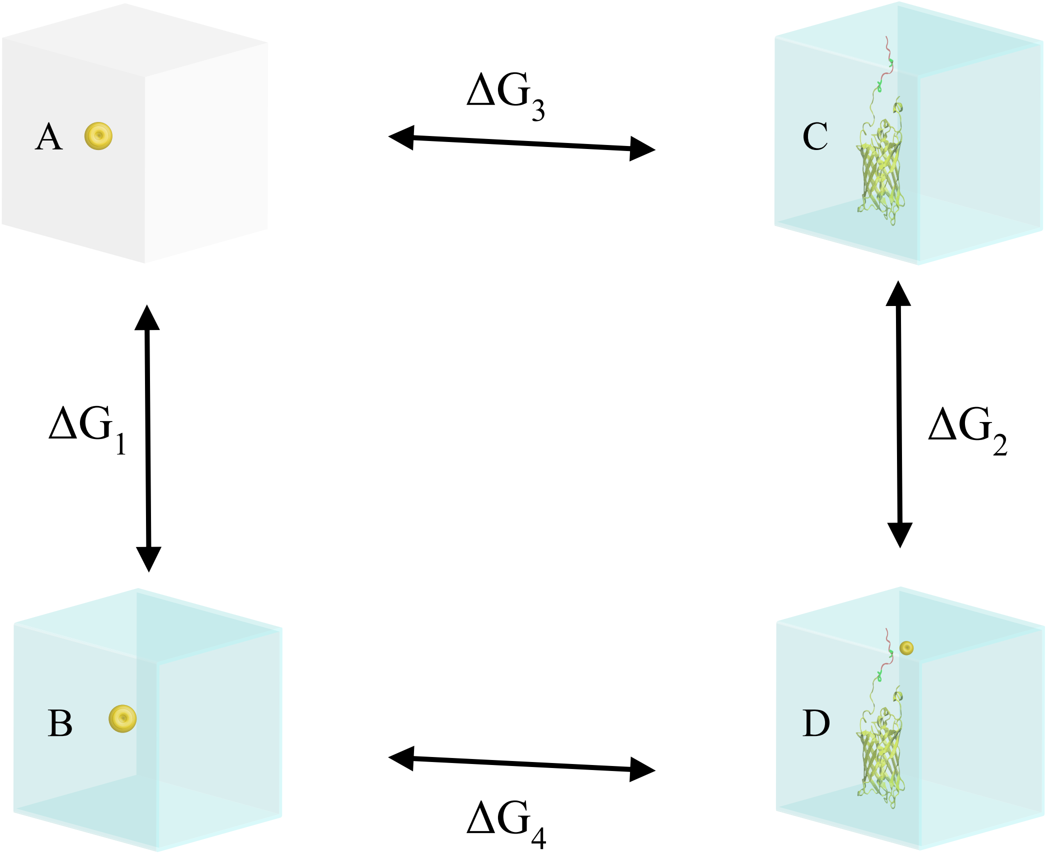


**Fig. S5.** Schematic representation of thermodynamics cycle for calculating the relative binding free energy of palladium binding to the histidine of peptides.(A-B) Solvation free energy of palladium atom in the aqueous solution. (C-D) Relative free energy binding of the palladium with the histidines of the peptides. overall binding free energy is derived from $\Delta G_{binding} = \Delta G_2 - \Delta G_1$

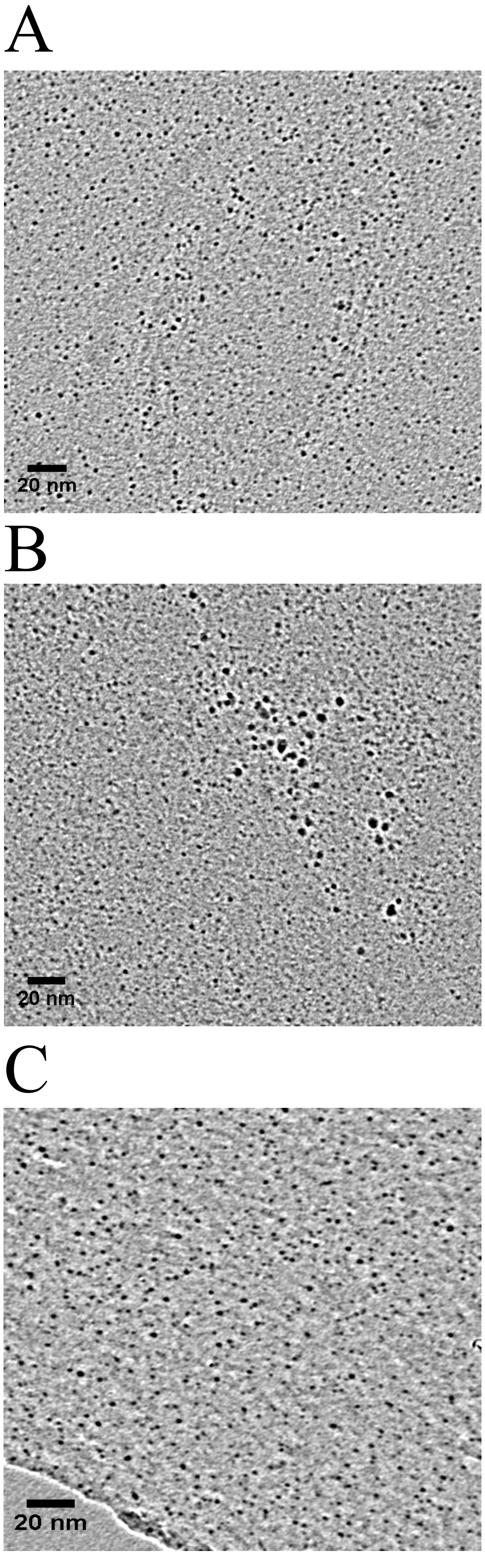


**Fig. S6.** (A-C) TEM images of nanoparticles produced from GFPuv-attached peptides Pd4 (A), A6 (B) & A11 (C).

**Table S1: Optimized condition via coupling reaction of iodobenzene and phenylboronic acid. Solvent, base, and temperature.**

$$\text{Ar–X}+Ar' \xrightarrow[Solvent,Time,andTemperature]{Catalyst,Base} Ar - Ar' \qquad (1)$$

| Entry | Solvent | Base (mmol) | Time (h) | Temperature (°C) | Yield[b] (%) |
|---|---|---|---|---|---|
| **1** | EtOH: $H_2O$ (1:1) | KOtBu | 3.0 | 80 | 47 |
| **2** | EtOH: $H_2O$ (1:1) | $K_2HPO_4$ | 3.0 | 80 | 65 |
| **3** | EtOH: $H_2O$ (1:1) | $K_2HPO_4$ | 3.0 | 80 | 62 |
| **4** | EtOH: $H_2O$ (1:1) | $K_2CO_3$ | 3.0 | 80 | 98 |
| **5** | EtOH: $H_2O$ (1:1) | $K_2CO_3$ | 8.0 | 80 | 82 |
| **6** | EtOH: $H_2O$ (3:1) | $K_2CO_3$ | 1.5 | 80 | 98 |
| **7** | EtOH: $H_2O$ (1:1) | $K_2CO_3$ | 1.5 | 80 | 97 |
| **8** | EtOH: $H_2O$ (1:3) | $K_2CO_3$ | 3.5 | 80 | 74 |

Reaction conditions:iodobenzene (0.1 mmol), phenylboronic acid (0.12 mmol), base (0.3 mmol) and Pd NPs (0.005 $\mu$mol) were mixed in solvent and refluxed under $N_2$. ([b])Yield were determined by HPLC.

**Table S2: Optimized condition via coupling reaction of iodobenzene and phenyltin trichloride. Solvent, base, and temperature.**

$$\text{Ar–X} + Ar - SnCl_3 \xrightarrow[EtOH:H\_2OandRefulx]{Catalyst,Base} Ar - Ar' \qquad (2)$$

| Entry | Solvent | Pd (mmol %) | Time (h) | Temperature (°C) | Yield[b] (%) |
|---|---|---|---|---|---|
| **1** | 5 | KOH | 6.0 | 80 | 72 |
| **2** | 5 | $K_2CO_3$ | 6.0 | 80 | 34 |
| **3** | 5 | $K_2HPO_4$ | 6.0 | 80 | 36 |
| **4** | 5 | CsF | 6.0 | 80 | 97 |
| **5** | 5 | $K_3PO_4$ | 48.0 | 40 | 58 |
| **6** | 5 | $K_3PO_4$ | 20.0 | 60 | 70 |
| **7** | 10 | $K_3PO_4$ | 6.0 | 80 | 96 |
| **8** | 2 | $K_3PO_4$ | 16 | 80 | 88 |
| **9** | 1 | $K_3PO_4$ | 16 | 80 | 64 |

Reaction conditions:iodobenzene (0.1 mmol), phenylboronic acid (0.12 mmol), base (0.3 mmol) and Pd NPs (0.005 $\mu$mol) were mixed in solvent and refluxed under $N_2$. ([b])Yield were determined by HPLC.
